# Interdomain interactions regulate the localization of a lipid transfer protein at ER-PM contact sites

**DOI:** 10.1101/2020.10.19.345074

**Authors:** Bishal Basak, Harini Krishnan, Padinjat Raghu

## Abstract

During phospholipase C-β (PLC-β) signalling in *Drosophila* photoreceptors, the phosphatidylinositol transfer protein (PITP) RDGB, is required for lipid transfer at endoplasmic reticulum (ER)-plasma membrane (PM) contact sites (MCS). Depletion of RDGB or its mis-localization away from the ER-PM MCS results in multiple defects in photoreceptor function. Previously, the interaction between the FFAT motif of RDGB and the integral ER protein dVAP-A was shown to be essential for accurate localization to ER-PM MCS. Here, we report that the FFAT/dVAP-A interaction alone is insufficient to localize RDGB accurately; this also requires the function of the C-terminal domains, DDHD and LNS2. Mutations in each of these domains results in mis-localization of RDGB leading to loss of function. While the LNS2 domain is necessary, it is not sufficient for the correct localization of RDGB, which also requires the C-terminal DDHD domain. The function of the DDHD domain is mediated through an intramolecular interaction with the LNS2 domain. Thus, interactions between the additional domains in a multi-domain PITP together lead to accurate localization at the MCS and signalling function.

## Introduction

The close approximation of intracellular membranes without fusion between them is emerging as a theme in cell biology (Gatta and Levine, 2017). Such apposition of membranes, referred to as membrane contact sites (MCS) can occur between multiple cellular organelles; most frequently, the endoplasmic reticulum (ER) which is the largest organelle, makes MCS with other cellular organelles including the plasma membrane (PM) (Cohen, Valm and Lippincott-Schwartz, 2018). ER-PM contact sites have been described in multiple eukaryotic cells, and are proposed to regulate a range of molecular processes including calcium influx and the exchange of lipids (Saheki and Camilli, 2017; Chen, Quintanilla and Liou, 2019).

The transfer of lipids between organelle membranes is a key function proposed for MCS. In the case of ER-PM contact sites, multiple lipids are thought to be transferred including phosphatidylserine (PS), phosphatidylinositol (PI), phosphatidic acid (PA), cholesterol and phosphatidylinositol 4-phosphate (PI4P) (Cockcroft and Raghu, 2018). These transfer activities are performed by several classes of lipid transfer proteins (LTPs). In order to carry out this function effectively, it is essential that these LTPs are accurately localized to ER-PM MCS, and several mechanisms that underlie this localization have been proposed (Alli-Balogun and Levine, 2019). LTPs frequently have multiple domains in addition to a lipid transfer domain. Some of these domains have been proposed to contribute to localization at the MCS but the *in vivo* function of several others is not clear. One group of LTPs named phosphatidylinositol transfer proteins (PITPs) mediate the specific transfer of PI between compartments. The first PITP identified and cloned was a protein with a single phosphatidylinositol transfer domain (PITPd) (Dickeson *et al*., 1989). Since then multiple PITPs, with either single or multiple domains have been identified in various species [reviewed in (Carvou *et al*., 2010)]. Importantly, in multi-domain PITPs, although the essential function of lipid transfer is conserved and restricted to the PITPd, the contribution of the additional domains to the regulation of PITPd activity *in vivo* is poorly understood.

*Drosophila* photoreceptors have emerged as an influential model system for the analysis of ER-PM contact sites (Yadav, Cockcroft and Raghu, 2016). Photoreceptors are polarized cells whose apical PM, also called rhabdomere, forms contact sites with the sub-microvillar cisternae (SMC), a specialized domain of the photoreceptor ER **[Figure 1A]**. The apical PM and the SMC are specialized to mediate sensory transduction through G-protein coupled Phospholipase C-β (PLC-β) activation (Raghu, Yadav and Mallampati, 2012). PLC-β activation triggers a series of enzymes whose substrates and products are lipid intermediates of the “PIP_2_ cycle” (Cockcroft and Raghu, 2016) that are distributed between the apical PM and the SMC. Some of these lipid intermediates such as PI and PA need to be transported between the apical PM and the SMC. *Drosophila* photoreceptors express a large multidomain protein, **R**etinal **D**e**g**eneration **B** (RDGB) that has a well-annotated PITPd (RDGB^PITPd^). Loss of function or hypomorphic mutants for *rdgB* represented by *rdgB*^*2*^ and *rdgB*^*9*^ alleles respectively, show defective electrical responses to light, retinal degeneration and defects in light activated PIP_2_ turnover. RDGB^PITPd^ has been shown to bind and transfer PI and PA *in vitro*, and is sufficient to support aspects of RDGB function *in vivo* (Yadav *et al*., 2015). Interestingly, the RDGB protein is localized exclusively to the MCS between the apical PM and the SMC (Vihtelic *et al*., 1993) **[Figure 1 A]**, thus offering an excellent *in vivo* setting to understand the relationship between LTP activity at an ER-PM contact site, and its physiological function. RDGB is a large multidomain protein; in addition to the N-terminal PITPd, the RDGB protein also includes several other domains including an FFAT motif, a DDHD domain and LNS2 domain **[Figure 1 B-**RDGB]. Of these, the interaction of the FFAT motif with the ER integral protein, dVAP-A has been shown to be important for the localization and function of RDGB *in vivo* (Yadav *et al*., 2018). However, the function of the two additional C-terminal domains: DDHD and LNS2 in the context of full length protein remain unknown. In cultured cells, the LNS2 domain of Nir2, the mammalian homologue of RDGB has been reported to have a role in localizing the protein to the PM (Kim *et al*., 2013, 2015) but the physiological significance of this is not known.

**Figure 1:**
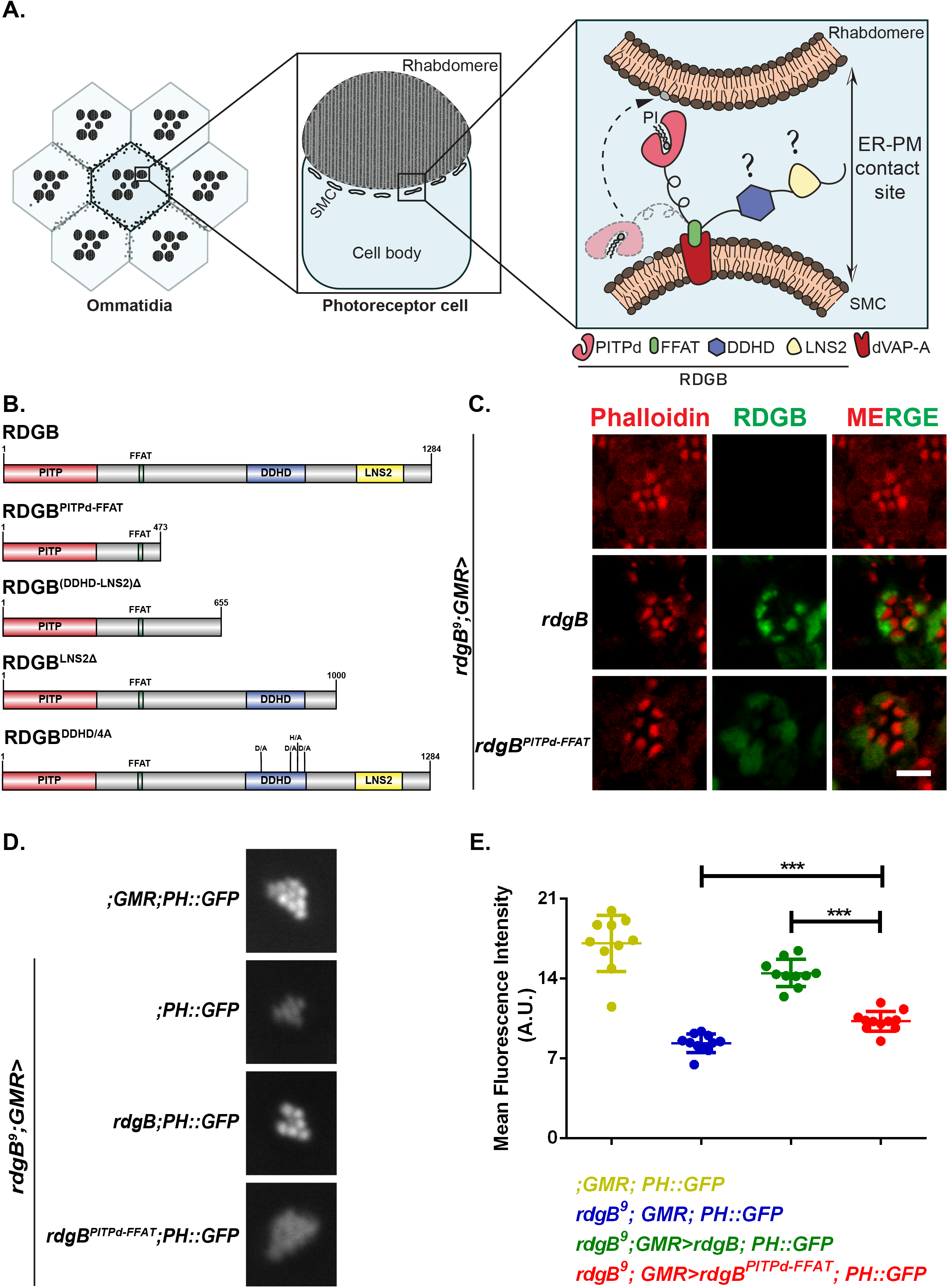
The PITPd and FFAT motif are insufficient for accurate localization of RDGB at ER-PM junctions in *Drosophila* photoreceptors. A. The *Drosophila* eye is composed of repeating units called the ommatidia each of which includes photoreceptor cells. Cross section of an individual photoreceptor is shown. The apical PM is thrown into numerous microvillar projections collectively termed as the rhabdomere, while the modified smooth ER compartment called Sub Microvillar Cisternae (SMC) is present at a distance of approximately ∼10 nm from it. Expanded view of a membrane contact site is depicted. Rhabdomere and SMC membranes are marked which form the ER-PM contact site. dVAP-A and RDGB protein with its individual domains are shown. Domains whose function is investigated here are marked with ?. Dotted arrow indicated the proposed movement of the PITPd to transfer phosphatidylinositol (PI) from the SMC to the PM. B. Domain structure of RDGB and the list of constructs generated in this study. RDGB protein is 1284 amino acid long and contains three domains-PITPd (red), DDHD (blue) and LNS2 (yellow), a FFAT motif (green). The length of the protein is marked on each of the construct. Individual deletion constructs of RDGB used in this manuscript are depicted. RDGB^DDHD/4A^ represents the full length RDGB where each of the conserved residues (D,D,H and D) in the DDHD domain has been mutated to alanine [Domain structure of RDGB drawn using Illustrator for Biological Sequences (IBS) software; http://ibs.biocuckoo.org/]. C. Confocal images of retinae obtained from flies of the mentioned genotypes. Transverse sections of an individual ommatidium are shown. Red represents phalloidin which marks the rhabdomeres and green represents immunostaining with an antibody to RDGB. Scale bar= 5 µm. D. Representative images of fluorescent deep pseudopupil from 1 day old flies of the mentioned genotypes expressing the PH-PLCδ::GFP probe. E. Quantification of the fluorescent deep pseudopupil (A.U.=Arbitrary Units). Y-axis denotes the mean intensity per unit area ±s.e.m.. Individual genotypes depicted are marked. n>=10 flies per genotype (***P<0.001, two tailed unpaired t-test).

The additional 180 amino acid long DDHD domain was first noted in Nir2 (Lev *et al*., 1999) and subsequently in the phosphatidic acid preferring phospholipase A1 (PLA1) family of proteins, first purified by Higgs & Glomset (Higgs and Glomset, 1994). This domain is named on the basis of 4 conserved amino acids D, D, H and D that are predicted to form a divalent metal binding site based on pattern of metal binding residues seen in phosphoesterase domains. In mammals, there are three members in Phosphatidic acid preferring phospholipase A1 family all of which possess the DDHD domain: PA-PLA_1_/DDHD1, KIAA072p/DDHD2 and p125/Sec23ip; mutations in DDHD2 have been found in patients with the neurodegenerative disease Hereditary spastic paraplegia (Pensato *et al*., 2014; Nicita *et al*., 2019) and those in DDHD1 with SPG28 (Tesson *et al*., 2012). However, the cellular mechanism through which mutations in DDHD1 and DDHD2 lead to neurodegeneration remain unknown. Studies done on DDHD2 have shown that the DDHD domain in association with a motif called sterile alpha-motif (SAM) binds PI4P (Inoue *et al*., 2012). This binding to PI4P has been shown to be essential for targeting this domain to Golgi and ERGIC compartments both of which are enriched in PI4P. Further, the first three D, D and H residues have been shown to be essential for the phospholipase activity of DDHD1 and KIAA072p. Another study (Klinkenberg *et al*., 2014) on the DDHD domain of p125/Sec23ip shows that the DDHD domain alone binds to weakly acidic lipids such as PA, PS, PIPs and PIP_2_s. The presence of a SAM motif along with the DDHD domain renders the specific binding to PIPs, PA and PS. However, the DDHD domain of p125 was also targeted to PI4P enriched Golgi membranes indicating the specificity of the DDHD domain to PI4P. However, to date there has been no study on the importance if any of the DDHD domain in the RDGB/Nir2 family of proteins for either localization or function.

In this study, we report that in *Drosophila* photoreceptors, the FFAT motif is insufficient for accurately localizing RDGB at the ER-PM MCS, and also requires the presence of the C-terminal domains, DDHD and LNS2. Loss of the LNS2 domain of RDGB leads to both mis-localization of the protein away from ER-PM contact sites as well as loss of function. Additionally, mutation of the four conserved residues of the DDHD domain also leads to both mis-localization and loss of RDGB function *in vivo*. Lastly, we find that the DDHD domain physically interacts with the LNS2 domain and this interaction influences localization. Thus we hypothesize that interdomain interactions in the RDGB protein are required for accurate localization of RDGB to ER-PM junctions, and hence function *in vivo*.

## Results

### The PITPd and FFAT motif of RDGB is insufficient for RDGB function at ER-PM contact sites

When the PITPd of RDGB is expressed in photoreceptors, it is distributed diffusely in the cell body. In addition, in the context of the full-length protein, the FFAT motif has been found to be important for localizing RDGB at the ER-PM junction (Yadav *et al*., 2018). Hence, we asked if expressing just the portion of RDGB that includes only the PITPd and the FFAT motif is sufficient to correctly localize the protein to ER-PM junctions. Towards this, we generated a truncated construct of RDGB removing everything C-terminal to the FFAT motif named as RDGB^PITPd-FFAT^ **[Figure 1 B-**RDGB^PITPd-FFAT^], and expressed it in *rdgB*^*9*^ photoreceptors **[Supplemental data 1A]**. We determined the localization of this protein by immunostaining with an antibody raised to the PITPd. Unlike full length RDGB which localized at the ER-PM junction, RDGB^PITPd-FFAT^ was found to be mislocalized from the base of the rhabdomere **[Figure 1 C]** and distributed throughout the cell body. This indicates that while the FFAT motif is essential, it is not sufficient for accurate localization of RDGB at the base of the rhabdomere. RDGB is essential to support the levels of PIP_2_ at the apical PM by transferring PI at the ER-PM junction (Yadav *et al*., 2015). We tested if RDGB^PITPd-FFAT^ could support the function of RDGB in supporting PIP_2_ levels at the apical PM. PIP_2_ levels at the apical PM were quantified through the fluorescence of PH-PLCδ::GFP probe in the pseudopupil of the eye (Chakrabarti *et al*., 2015). As previously reported we found that the resting level of PIP_2_ at the apical PM of *rdgB*^*9*^ was reduced and could be restored to wild type levels by reconstitution with a wild type RDGB transgene (Yadav *et al*., 2015). When tested for the ability to rescue the reduced PIP_2_ levels in *rdgB*^*9*^ flies, RDGB^PITPd-FFAT^ was found to rescue the defect only partially. As compared to *rdgB*^*9*^ flies, PIP_2_ levels were found to be higher in *rdgB*^*9*^; *GMR>rdgB*^*PITPd-FFAT*^ than in *rdgB*^*9*^ but was significantly lower than that of the wild type controls **[Figure 1 D, E]**. The expression levels of the PH-PLCδ::GFP probe were found to be similar across all genotypes, implying that the reduced fluorescence was a direct read out of the reduced PIP_2_ levels at the ER-PM MCS **[Supplemental data 1B]**. Collectively, these results imply that the domains present C-terminus to the FFAT motif contribute to localizing RDGB correctly which then impact its function.

### Loss of LNS2 domain from RDGB leads to loss of *in vivo* function

There are two well annotated domains C-terminal to the FFAT motif in RDGB: DDHD and LNS2. Of these, the LNS2 domain has previously been implicated in the membrane localization of Nir2, the mammalian orthologue of RDGB (Kim *et al*., 2013, 2015). To understand if the C-terminal domains are essential for localization and function of RDGB, we removed the C-terminal of the protein from just before the start of the DDHD domain **[Figure 1 B**-RDGB^(DDHD-LNS2)Δ^] and expressed this protein in *rdgB*^*9*^ photoreceptors [*rdgB*^*9*^; GMR>*rdgB*^(DDHD-LNS2)Δ^] **[Supplemental data 2A]**. Immunolocalization experiments revealed that RDGB^(DDHD-LNS2)Δ^ was not localized to the ER-PM contact site but was distributed throughout the cell body **[Figure 2 A]**. An important physiological output of phototransduction is the generation of an electrical response to light; this is typically measured using an electroretinogram (ERG) and the amplitude of the ERG is reduced in *rdgB* mutants. Further, we found that RDGB^(DDHD-LNS2)Δ^ was unable to rescue the ERG phenotype of *rdgB*^*9*^ **[Figure 2 B, Supplemental data 2B]** and the PIP_2_ levels in *rdgB*^*9*^ flies expressing RDGB^(DDHD-LNS2)Δ^ were comparable to that of *rdgB*^*9*^ flies **[Figure 2 C, Supplemental data 2C]**, although probe levels were found to be unaltered across all genotypes **[Supplemental data 2D]**. These findings imply that the presence of one or both of these domains is essential for correct localization and function of RDGB.

**Figure 2:**
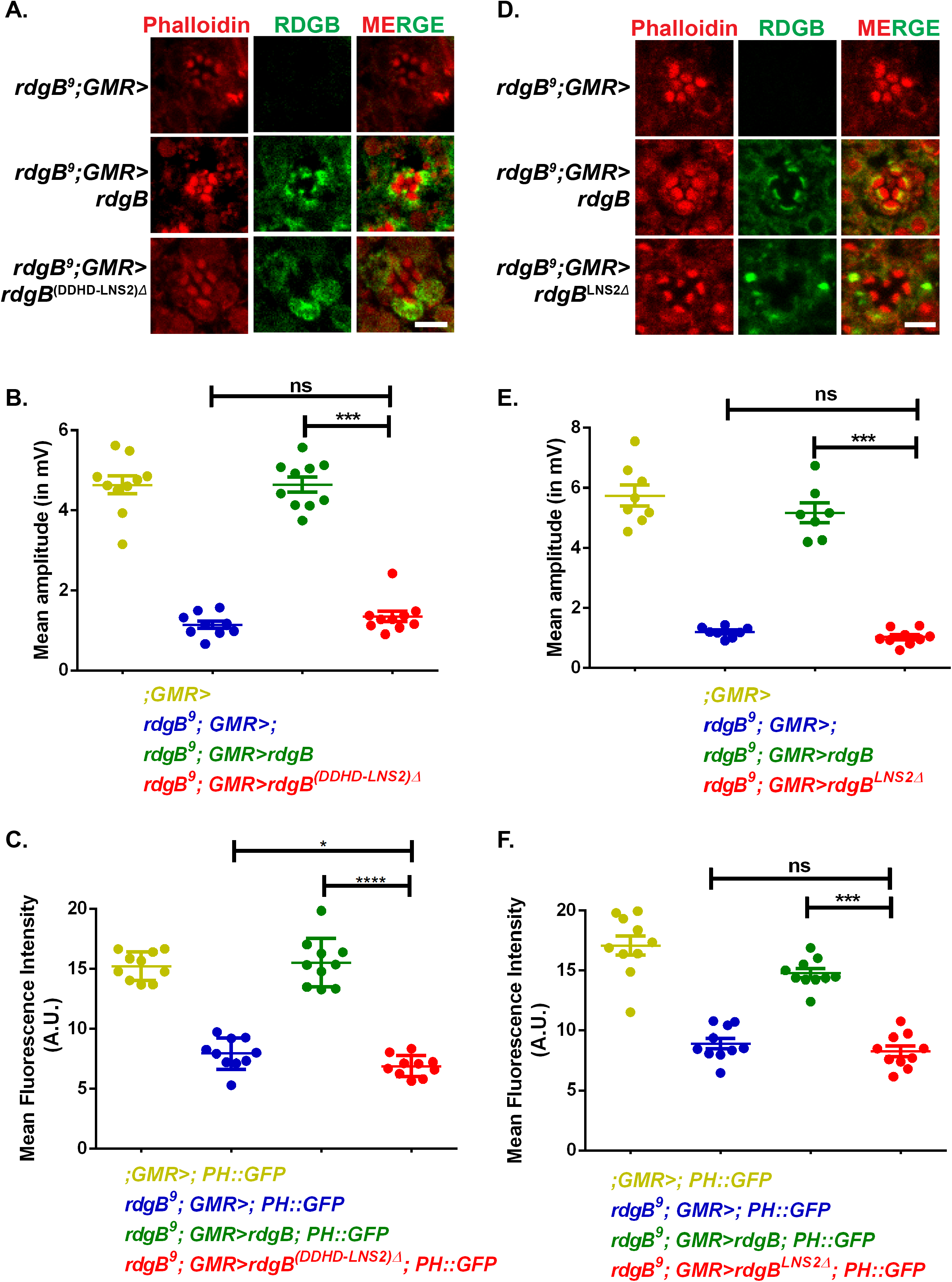
The DDHD and LNS2 domains of RDGB are indispensable to support full RDGB function. A. Confocal images of retinae obtained from flies expressing RDGB^(DDHD-LNS2)Δ^ and controls. Transverse sections of an individual ommatidium are shown. Red represents phalloidin which marks the rhabdomeres and green represents immunostaining for the RDGB protein. Scale bar= 5 µm. B. Quantification of the light response from 1 day old flies of RDGB^(DDHD-LNS2)Δ^ and controls. Y-axis represents mean amplitude (mV) ±s.e.m. n=10 flies per genotype (***P<0.001, ns= not significant; two tailed unpaired t-test). C. Quantification of the fluorescence intensity of the deep pseudopupil of RDGB^(DDHD-LNS2)Δ^ and controls. (A.U. =Arbitrary Units). n=10 flies per genotype. Y-axis denotes the mean intensity per unit area ±s.e.m. (***P<0.001, two tailed unpaired t-test). D. Confocal images of retinae obtained from flies expressing RDGB^LNS2Δ^ and controls. Transverse sections of an individual ommatidium are shown. Red represents phalloidin which marks the rhabdomeres and green represents immunostaining for RDGB. Scale bar= 5 µm. E. Quantification of the light response from 1 day old flies expressing RDGB^LNS2Δ^ and controls. Each point on Y-axis represents mean amplitude ±s.e.m., n>=7 flies per genotype (*** - P<0.001, ns= not significant; two tailed unpaired t-test). F. Quantification of the fluorescence intensity of the deep pseudopupil from 1 day old flies expressing RDGB^LNS2Δ^ and controls. Y-axis denotes the mean intensity per unit area (A.U. =Arbitrary Units) ±s.e.m., n=10 flies per genotype (*** P<0.001, two tailed unpaired t-test).

Since our data shows that loss of both domains together lead to complete loss of RDGB function we then went onto investigate the role played by each of these individual domains. Firstly, to test if the LNS2 domain in RDGB is required for localization, we deleted the LNS2 domain **[Figure 1 B**-RDGB^LNS2Δ^] and expressed the rest of the RDGB protein in photoreceptors of *rdgB*^*9*^ flies (*rdgB*^*9*^; GMR>*rdgB*^LNS2Δ^) **[Supplementary data 2E]**. RDGB^LNS2Δ^ was found to be completely mislocalized from the base of the rhabdomere, suggesting that this domain is indispensable for localization of RDGB **[Figure 2 D]**. We then performed ERG recordings to test if the LNS2 domain has a role in supporting RDGB function *in vivo*. We found that the electrical response to light measured in RDGB^LNS2Δ^ expressing photoreceptors was as low as that in *rdgB*^*9*^ **[Figure 2 E, Supplemental data 2F]**. Similarly, PM PIP_2_ levels in *rdgB*^*9*^ reconstituted with RDGB^LNS2Δ^ (*rdgB*^*9*^; *GMR>rdgB*^LNS2Δ^) was found to be as low as in *rdgB*^*9*^ photoreceptors **[Figure 2 F, Supplemental data 2G]** although probe levels were equal across all genotypes **[Supplemental data 2H]**. These results collectively support an indispensable role for the LNS2 domain in supporting RDGB localization and function *in vivo*.

### The LNS2 domain is an apical PM binding signal in RDGB

Our *in vivo* analysis reveals that loss of LNS2 domain severely affects RDGB localization and function at ER-PM MCS. While the integral ER membrane protein dVAP-A has been previously implicated in localizing RDGB to the MCS by interacting with the latter’s FFAT motif, we questioned what additional factors might be contributing for accurate localization of RDGB at the ER-PM MCS. For this we developed a sub cellular fractionation assay and found that in *Drosophila* photoreceptors, RDGB is a membrane associated protein which co-fractionates with the membrane marker, dVAP-A **[Figure 3 A, A’]**. However, when the LNS2 domain is deleted from RDGB, the protein RDGB^LNS2Δ^ now mainly co-fractionates with the cytosolic protein tubulin. This implies that the LNS2 domain is essential for membrane association of RDGB and its loss from the protein makes RDGB cytosolic **[Figure 3 B, B’]**.

**Figure 3:**
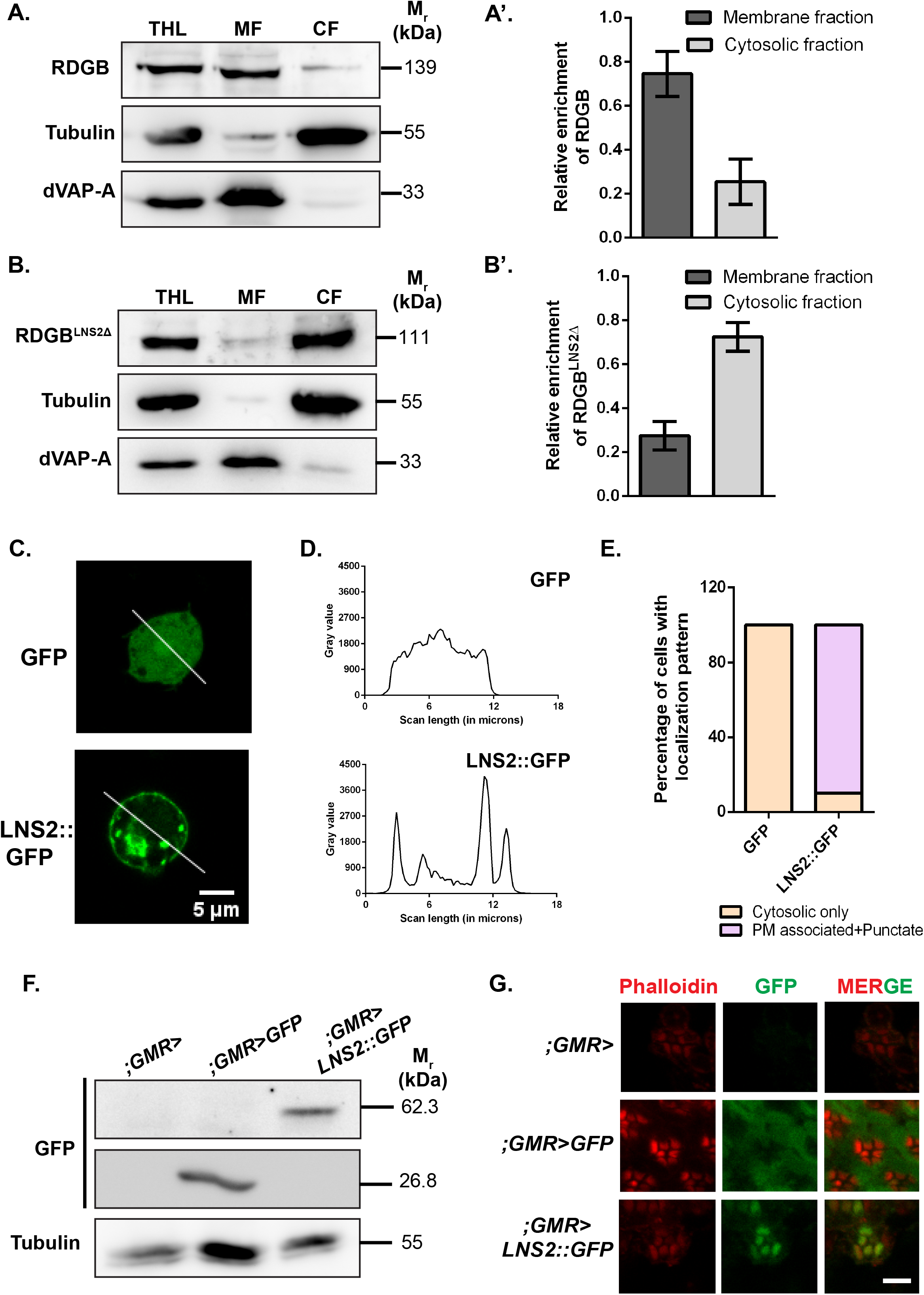
The LNS2 domain is an apical PM targeting signal. A. Representative immunoblot showing fractionation of RDGB between the membrane and cytosolic fractions from *Drosophila* heads. dVAP-A, an ER integral protein marks the membrane fraction, while the soluble protein tubulin represents the cytosolic fraction [THL=Total Head Lysate, MF=Membrane fraction, CF=Cytosolic fraction] (N=3). A’. Quantification showing the relative enrichment of RDGB in membrane and cytosolic fractions. The Y-axis, denoting the relative enrichment, is calculated as the ratio of RDGB in each fraction to the sum total of RDGB in both membrane and cytosolic fractions. B. Representative immunoblot showing fractionation of RDGB^LNS2Δ^ between the membrane and cytosolic fractions from *Drosophila* heads. dVAP-A, an ER integral protein represents the membrane fraction, while the soluble protein tubulin represents the cytosolic fraction. [THL=Total Head Lysate, MF=Membrane fraction, CF=Cytosolic fraction] (N=3). B’. Quantification showing the relative enrichment of RDGB^LNS2Δ^ in membrane and cytosolic fractions. The Y-axis, denoting the relative enrichment, is calculated as the ratio of RDGB^LNS2Δ^ in each fraction to the sum total of RDGB^LNS2Δ^ in both membrane and cytosolic fractions. C. Confocal images of S2R+ cells transfected with pJFRC-GFP or pJFRC-LNS2::GFP. Green represents signal from GFP. The white line indicated the region of the cells selected for the line scan quantified in D. D. Line scan profiles showing the fluorescence intensity of GFP distributed along the line marked in C. Y-axis is the intensity of fluorescence while X-axis represents the length of the cell in µm. In GFP transfected cells, the fluorescence is distributed uniformly along the width of the cell while in LNS2::GFP, the highest intensity is seen at the PM and in punctate structures in the cytosol. E. Bar graph showing the distribution of localization patterns of GFP and LNS2::GFP in S2R+ cells (N= 30 cells). Y-axis indicates the proportion of cells showing either cytosolic or membrane associated pattern. F. Western blot of protein extracts made from 1 day old fly heads of the mentioned genotypes. The blot is probed with antibody to GFP. Tubulin is used as a loading control (N=3). G. Confocal images of retinae obtained from flies expressing LNS2::GFP, GFP or controls. Transverse sections of an individual ommatidium are shown. Red represents phalloidin which marks the rhabdomeres and green represents immunostaining for GFP. Scale bar= 5 µm.

While our sub-cellular fractionation assay reveals that the LNS2 domain is essential for membrane association of RDGB, it does not identify the cellular membrane to which the domain is targeted. To understand this, we cloned the LNS2 domain alone, tagged to GFP (LNS2::GFP) and expressed it in S2R+ cells. Under these conditions, LNS2::GFP was found to localize primarily to the PM with some punctate structures within the cell **[Figure 3 C, D, E]**. To test if the LNS2 domain is also able to localize to the PM in photoreceptors, we expressed LNS2::GFP in wild type photoreceptors **[Figure 3 F]**. Unlike GFP which showed a completely diffuse distribution in the photoreceptor cell body, LNS2::GFP was found to be localized very specifically to the rhabdomeres, i.e. the apical PM **[Figure 3 G]**. It is important to note that the photoreceptors of *Drosophila* are highly polarized cells and exhibit strikingly structural differences in the arrangement of its apical vs basolateral PM. While the LNS2 domain associates to the PM in unpolarised S2R+ cells, (similar to what has been reported for the LNS2 domain of Nir2), it localizes exclusively to the apical PM and not the basolateral PM in polarized photoreceptor cells implying underlying mechanisms which allow this preferential binding.

### The DDHD domain is required for normal localization and function of RDGB

If the FFAT motif is essential for interaction with the ER (via dVAP-A) and the LNS2 domain with the apical PM at the ER-PM MCS of *Drosophila* photoreceptors, then what is the function of the DDHD domain, present just N-terminal to the LNS2 domain? To determine if this domain is essential for the function of RDGB, we at first checked if at all the residues which give the domain its identity and nomenclature are present in RDGB. For this, we aligned the DDHD domain of PA-PLA1 with that of RDGB and determined that all four residues D, D, H and D are indeed also conserved in RDGB **[Figure 4 A]**. To check if these conserved residues are functionally important we mutated these 4 residues each to alanine **[Figure 1 B**-RDGB^DDHD/4A^] in the full length protein, expressed it in fly photoreceptors **[Supplemental data 3A]** and checked for its localization. RDGB^DDHD/4A^ was found to be diffusely distributed and not localized to the base of the rhabdomere **[Figure 4 B]**. Thus the conserved residues of the DDHD domain are essential to localize RDGB to ER-PM MCS.

**Figure 4:**
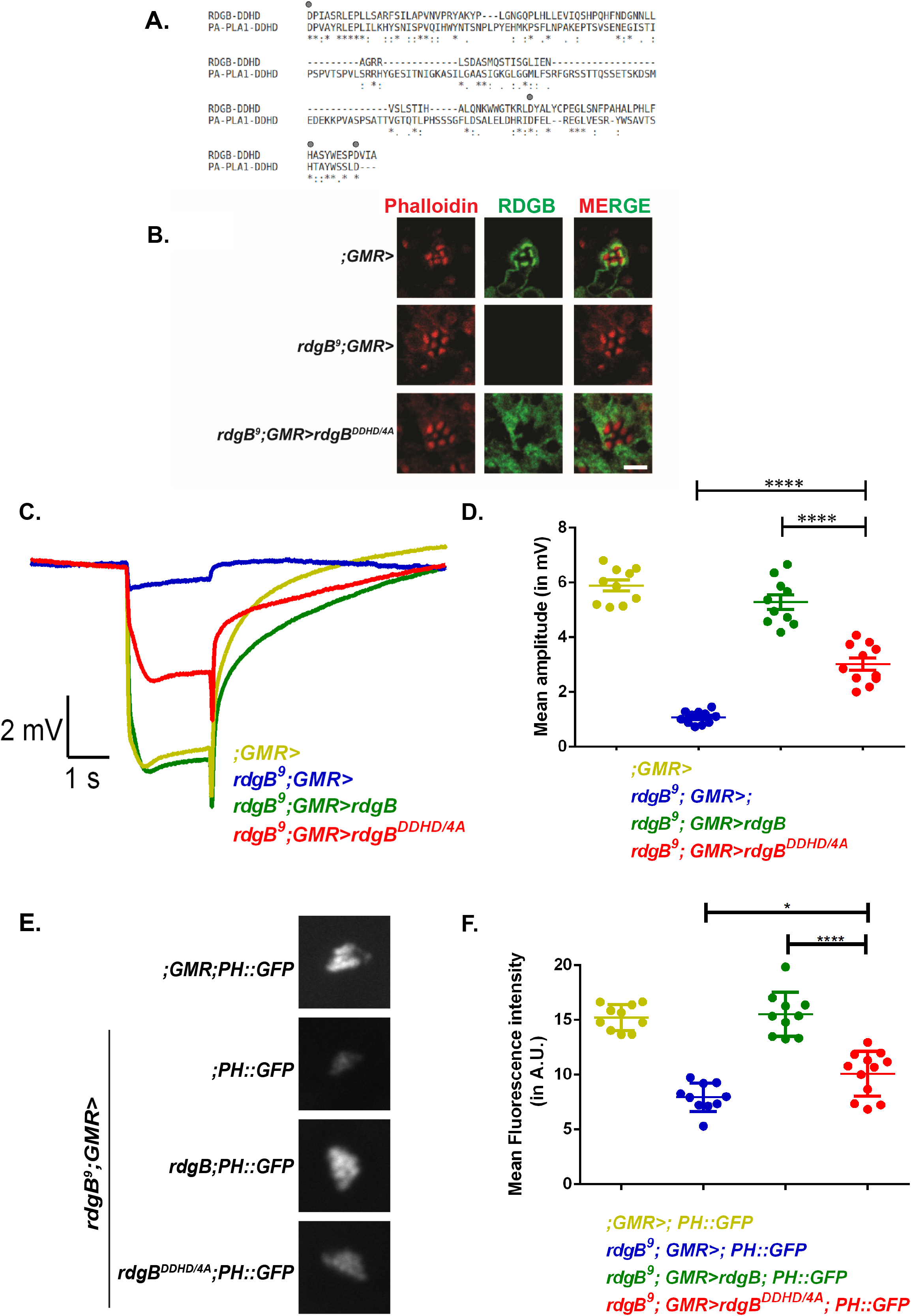
The 4 conserved residues (D, D, H and D) of the DDHD domain are essential to support RDGB function *in vivo*. A. Alignment of DDHD domain region of RDGB protein with that from the DDHD1/PA-PLA1 protein. Residues 776 to 905 of RDGB protein are aligned to residues 669 to 854 of PA-PLA1 using ClustalO. ‘:’ indicates that one of the following ‘strong’ groups is fully conserved:-STA, NEQK, NHQK, NDEQ, QHRK, MILV, MILF, HY, FYW. ‘.’ indicates that one of the following ‘weaker’ groups is fully conserved:-CSA, ATV, SAG, STNK, STPA, SGND, SNDEQK, NDEQHK, NEQHRK, FVLIM, HFY. The four residues D, D, H and D which are considered functionally important to this domain are marked with a grey circle on the alignment. B. Confocal images of retinae obtained from flies expressing RDGB^DDHD/4A^ and controls. Transverse sections of an individual ommatidium are shown. Red represents phalloidin which marks the rhabdomeres and green represents immunostaining for the RDGB protein. Scale bar= 5 µm. C. Representative ERG traces from 1 day old flies expressing RDGB^DDHD/4A^ and the relevant controls. Y-axis represents ERG amplitude in mV, X-axis represents time in sec. Genotypes studied are indicated. D. Quantification of the light response from 1 day old flies expressing RDGB^DDHD/4A^ and controls. Each point on Y-axis represents mean amplitude ±s.e.m., n>=10 flies per genotype (*** - p<0.001, ns= not significant; two tailed unpaired t-test). E. Representative images of fluorescent deep pseudopupil from 1 day old flies expressing RDGB^DDHD/4A^ and controls expressing the PH-PLCδ::GFP probe. F. Quantification of the fluorescence intensity of the deep pseudopupil from flies expressing RDGB^DDHD/4A^ and controls. Y-axis denotes the mean intensity per unit area (A.U. =Arbitrary Units) ±s.e.m., n>=10 flies per genotype (*** P<0.001, two tailed unpaired t-test).

Since altered localization leads to defects in RDGB function, we then tested if mutation of these conserved residues in the full length protein also had a similar impact. We found the electrical response to light in *rdgB*^*9*^ photoreceptors expressing RDGB^DDHD/4A^ was significantly lower than that of wild type **[Figures 4 C, D]**. Similarly, we found that the PIP_2_ levels in *rdgB*^*9*^ photoreceptors reconstituted with RDGB^DDHD/4A^ were as low as in *rdgB*^*9*^ **[Figures 4 E**, **F]**, although probe levels were equivalent in all genotypes **[Supplemental data 3B]**. These results collectively suggest that the DDHD domain is required for the correct localization and normal function of RDGB.

### The DDHD domain interacts with the LNS2 domain

Our *in vivo* data shows that mutations in the conserved residues of the DDHD domain impact localization and function of the full length protein. To understand the function of the DDHD domain as a whole, we expressed an mCherry tagged version of the DDHD domain in S2R+ cells. We found that DDHD domain showed a diffuse distribution in the majority of cells, while in some cells a few punctate structures were also observed **[Figure 5 A, B, C]**. Since there are now two individual domains, each of which when mutated leads to altered localization and loss of function, how do they contribute to the localization of RDGB? To analyze this, we generated an mCherry::DDHD-LNS2 construct and expressed it in S2R+ cells. In sharp contrast to the diffuse localization of the DDHD domain, mCherry::DDHD-LNS2 was found to have a punctate distribution very close to the PM. **[Figure 5 D, E, F]**. Likewise, the primarily PM localization of the isolated LNS2 domain was also altered. These findings suggest that the DDHD domain can modulate the localization of the LNS2 domain when present in *cis*.

**Figure 5:**
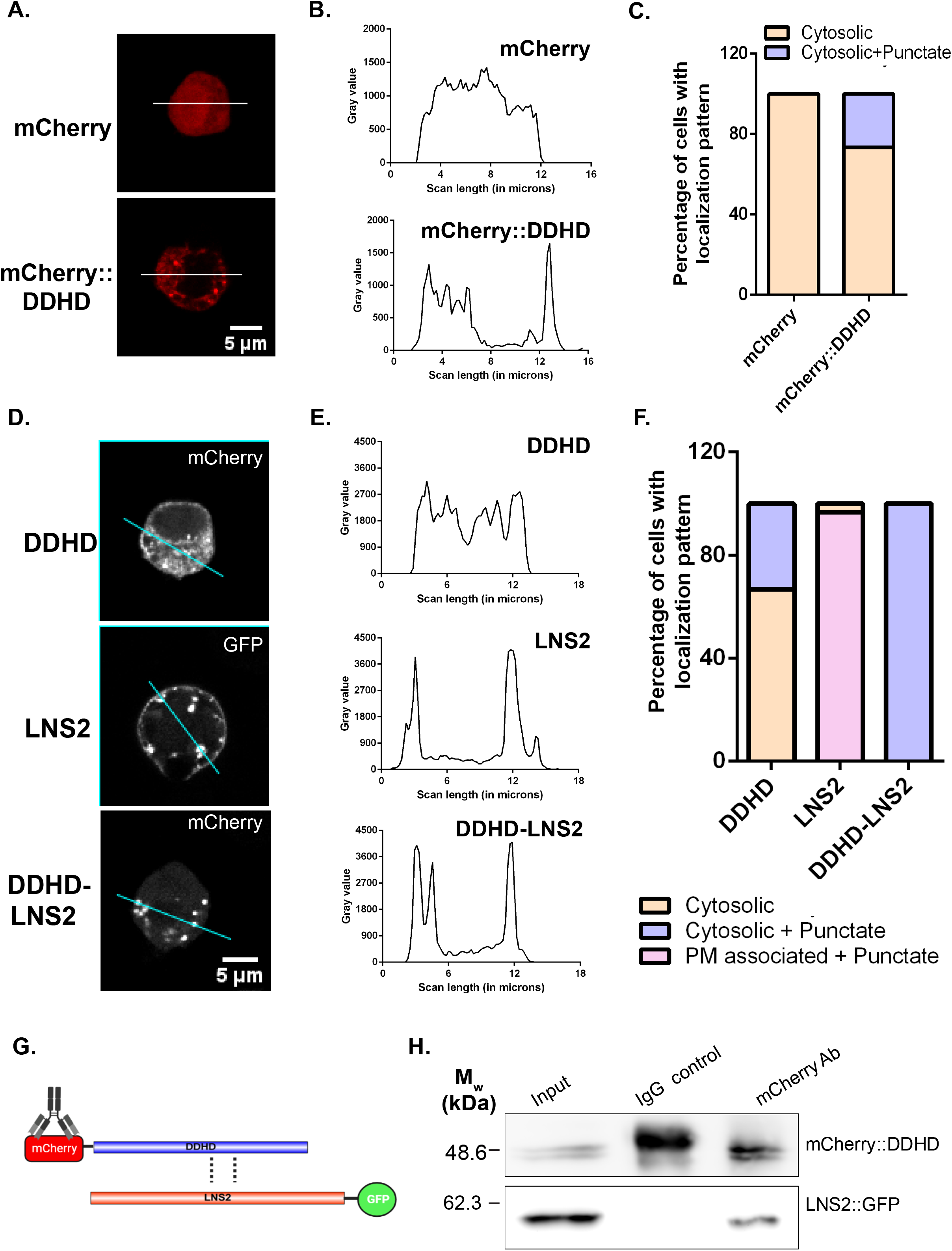
The DDHD domain physically interacts with the LNS2 domain to regulate the latter’s localization. A. Confocal images of S2R+ cells transfected with pUAST-mCherry or pUAST-mCherry::DDHD. Red represents mCherry. The white line indicated the region of the cells selected for the line scan quantified in B. B. Line scan profiles showing the fluorescence intensity of mCherry distributed along the line marked in A. Y-axis is the intensity of fluorescence while X-axis represents the length of the cell in µm. mCherry is distributed uniformly along the line in A for mCherry and mCherry::DDHD. C. Bar graph showing the distribution of localization patterns of mCherry and mCherry::DDHD in S2R+ cells (N= 30 cells). Y-axis indicates the proportion of cells showing either cytosolic or membrane associated pattern. D. Confocal images of S2R+ cells transfected with LNS2::GFP, mCherry::DDHD and mCherry::DDHD-LNS2. The cyan lines represent the regions of the cells selected for line scan in E. E. Line scan profiles showing the fluorescence intensity of mCherry or GFP distributed along the line marked in D. Y-axis is the intensity of fluorescence while X-axis represents the length of the cell in µm. The fluorescence intensity is distributed uniformly along the line in D for mCherry::DDHD, while it peaks at the PM and punctate structures for LNS2::GFP, and only at punctate structures in mCherry::DDHD-LNS2. F. Bar graph showing the distribution of localization patterns of mCherry::DDHD, LNS2::GFP and mCherry::DDHD-LNS2 in S2R+ cells (N= 30 cells). Y-axis indicates the proportion of cells showing either cytosolic or membrane associated pattern. G. Cartoon representing co-immunoprecipitation performed to test the interaction of DDHD domain with the LNS2 domain. Tags used for the individual protein domains are shown. Antibody used for the immunoprecipitation is indicated. Potential interactions being probed are shown in dotted lines. H. Representative immunoblot showing the co-immunoprecipitation of LNS2::GFP with mCherry::DDHD from S2R+ cells transfected with this combination of constructs. IgG control-negative control for immunoprecipitation. [Illustrations made using BioRender (https://biorender.com/) and Illustrator for Biological Sequences (IBS) (http://ibs.biocuckoo.org/)] (N=3).

One of the possible ways via which the DDHD domain can modulate the localization of the LNS2 domain is via physical interaction. To understand if indeed this is true, we co-expressed mCherry tagged DDHD domain (mCherry::DDHD) in S2R+ cells along with GFP tagged LNS2 domain (LNS2::GFP). When we immunoprecipitated the DDHD domain using an mCherry antibody, we could detect the LNS2 domain in the pulled down fraction implying physical interaction between these two domains **[Figure 5 G, H]**.

## Discussion

The presence of multiple domains in LTPs is hypothesized to enable their correct localization at MCS. These domains are conceptualized as independent units each with a unique property contributing to optimal lipid transfer function at MCS. A similar model has been proposed for the PITPs, a specific group of LTPs that can transfer PI at ER-PM junctions (Kim *et al*., 2013, 2015). However, in the case of *Drosophila* RDGB, a multidomain PITP, it has been noted that re-expression of just RDGB^PITPd^ which performs lipid transfer *in vitro*, in a null mutant background, is sufficient to rescue key phenotypes *in vivo* suggesting the sufficiency of the RDGB^PITPd^ in supporting RDGB function. A more recent study has however shown that while RDGB^PITPd^ can rescue key phenotypes, it is incapable of supporting lipid turn over during high rates of PLC-β signalling (Yadav *et al*., 2018), emphasizing the importance of ensuring a sufficiently high concentration of RDGB at the ER-PM contact site in photoreceptors **[Figure 6 A, B]**.

**Figure 6:**
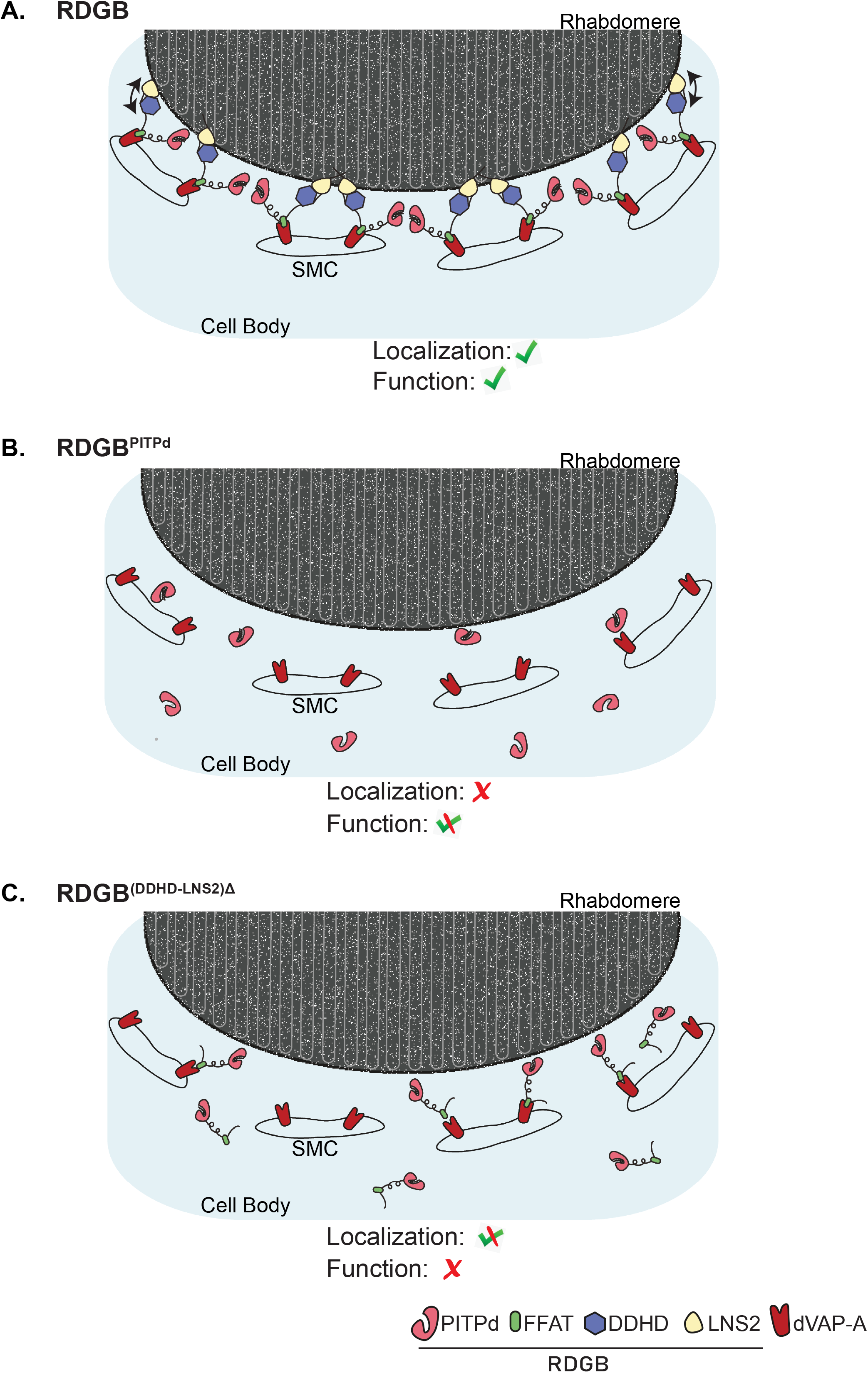
Model depicting mechanisms localizing RDGB to ER-PM MCS in photoreceptors. Cartoon depicting a cross-sectional view of a *Drosophila* photoreceptor with the apical plasma membrane (rhabdomere) and the sub-microvillar cisternae (SMC) forming a contact site; the cell body is also shown. A. Wild type RDGB interacts with the ER (via the FFAT-VAP interaction) and the PM (via the LNS2 domain) for accurate localization of the protein at the MCS. The arrow indicates the interaction of the DDHD with the LNS2 domain which contributes to localization. These interactions ensure that a high concentration of RDGB is present at the ER-PM contact site to mediate lipid transfer function. B. RDGB^PITPd^ cannot interact with the ER or PM. The soluble protein is able to diffuse throughout the cytosol both near the ER-PM MCS but also elsewhere in the cell body. Hence a lower concentration of PITPd is found near the MCS and can partially substitute for full length RDGB function. C. RDGB^(DDHD-LNS2)Δ^ cannot interact with the PM component of the MCS, and is hence mislocalized. The loss of the C-terminal domains: DDHD and LNS2 lead to complete loss of RDGB function, implying the requirement of these domains in full length context.

How is RDGB accurately localized such that it can be concentrated at the ER-PM MCS? It has previously been demonstrated (Yadav *et al*., 2018) that an interaction between the FFAT motif and dVAP-A is essential for the normal localization and function of RDGB. In this study, surprisingly, we found that an RDGB protein with only the PITPd (for function) and the FFAT motif (for ER anchoring) was (i) mislocalized away from the base of the rhabdomere and (ii) unable to restore RDGB function. These observations imply that additional regions of the RDGB protein, C-terminal to the FFAT motif are functionally important. To the C-terminus of the FFAT motif lies the DDHD and LNS2 domains. We observed that loss of these domains together from full length RDGB led to mislocalization and complete loss of function **[Figure 6 C]**. Additionally, our findings that mutation of the DDHD domain or loss of the LNS2 domain, completely mislocalizes RDGB away from the base of the rhabdomeres and also abrogates RDGB function support a role for each of these domains individually in the localization and function of RDGB. The LNS2 domain when expressed by itself localized to the PM in cultured *Drosophila* cells and specifically to the apical PM in photoreceptors. These data strongly support the function of the LNS2 domain as a PM localization signal. Although previous studies have implicated the LNS2 domain of Nir2, the mammalian ortholog of RDGB, in localization to the PM (Kim *et al*., 2013, 2015), our data are the first demonstration of the requirement of this domain in supporting physiological function *in vivo*. Interestingly, when expressed in photoreceptors, the LNS2 domain localized only to the apical PM (and not the basolateral PM) suggesting a unique apical domain interaction partner that localizes it here. Studies on Nir2 have suggested the LNS2 domain binds PA (Kim *et al*., 2013); while we also found that the LNS2 domain of RDGB also binds PA and PS **[Supplemental data 4A]**, neither of these lipids is unique to or enriched in the apical PM. Thus the signal through which the LNS2 domain interacts specifically with the apical PM remains to be determined.

If the FFAT motif of RDGB mediates its interaction with dVAP-A and the LNS2 domain with the PM, what role does the DDHD domain serve in the protein? Although the DDHD domain was first reported in Nir2 (Lev *et al*., 1999), its function in this protein has not been described. However, studies of mammalian PA-PLA1 have implicated the DDHD domain in localization and function (Inoue *et al*., 2012; Klinkenberg *et al*., 2014) but the mechanism has not been discovered. Our finding that mutation of the D, D, H and D residues of this domain to 4A in full length RDGB led to mis-localization support a role for this domain in the correct localization of RDGB. Surprisingly, and in sharp contrast to the LNS2 domain, when expressed by itself, the DDHD domain did not localize to the PM but showed a diffuse cytosolic distribution **[Figure 5 A, B, C]**. Thus, while the DDHD domain is essential for PM localization of RDGB, this domain in itself is not sufficient and cannot act as a primary membrane targeting signal. Interestingly, we found that when co-expressed with the LNS2 domain, the DDHD domain was able to alter the localization of the LNS2 domain and in immunoprecipitation experiments, the DDHD and LNS2 domains were able to physically interact **[Figure 5 H]**. These two findings strongly suggest that the DDHD domain is able to influence the function of the LNS2 domain and it is likely that through this mechanism it influences the localization of RDGB, rather than a direct role in membrane localization **[Figure 6 A, C]**. Interestingly, in the case of mammalian DDHD2, the DDHD domain appears to act in conjunction with the adjacent SAM motif (Inoue *et al*., 2012). It is noteworthy that the DDHD domain in RDGB interacts with and influences the localization of the LNS2 domain, a domain that binds PA (this study); this has also been shown for the LNS2 domain of Nir2 (Kim *et al*., 2013). Interestingly the only other known DDHD domain containing proteins are the family of PA preferring phospholipase A1 enzymes; the significance of this observation is unknown but may reflect the importance of DDHD domains in some classes of PA binding proteins. The molecular mechanism by which the DDHD domain influences the function of the LNS2 domain in localizing RDGB to MCS remains to be determined. However our findings on the role of a wild type DDHD domain in preventing retinal degeneration provide an insight into the cellular mechanisms that could explain the neurodegenerative phenotype seen in spastic paraplegias, in patients carrying mutations in human DDHD1 and DDHD2.

In summary, our study identifies the C-terminal domains of RDGB that play a key role in its localization and hence function. We define a novel intramolecular interaction between these domains that is required to facilitate accurate localization of RDGB at ER-PM contact sites. More generally, our study provides a framework for understanding the localization of multidomain PITPs at MCS and their function *in vivo*.

## Acknowledgements

This work was supported by the Department of Atomic Energy, Government of India, under project no. 12-R&D-TFR-5.04-08002 and 12-R&D-TFR-5.04-0900, a Wellcome-DBT India Alliance Senior Fellowship (IA/S/14/2/501540) to PR. We thank the Transgenic Fly Facility and Central Imaging Facility at NCBS for support. We thank Dr. Girish Ratnaparkhi from IISER Pune for providing us with the dVAP-A antibody.

## Materials and methods

### Fly stocks

All fly stocks were maintained at 25°C incubators with no internal illumination. Flies were raised on standard corn meal media containing 1.5% yeast. UAS-Gal4 system was used to drive expression in the transgenic flies.

### Molecular Biology

BDGP gold clone 09970 containing the *rdgB*-RA transcript was used as the parent vector for making various constructs of RDGB used for the experiments. The cDNA coding region corresponding to RDGB^PITPd-FFAT^ (amino acids 1-472) was subcloned into pUAST-attB by using the restriction enzymes *Not*I and *Xba*I (NEB). Similarly, for making RDGB^(DDHD-LNS2)Δ^ the cDNA corresponding to amino acids 1-655 was amplified, and for RDGB^LNS2Δ^ the cDNA corresponding to amino acids 1-1000 was amplified and then individually subcloned in *Not*I and *Xba*I digested pUAST-attB. For cloning of RDGB^DDHD/4A^, mutations were introduced in the *rdgB* cDNA corresponding to amino acid numbers 776, 872, 894 and 902. The resulting mutant gene *rdgB*^*DDHD/4A*^ where the 4 residues were substituted to alanine was then subcloned *in Not*I and *Xba*I digested pUAST-attB. To clone the LNS2 domain alone, the cDNA of RDGB corresponding to amino acids 947-1259 was subcloned in pJFRC::GFP vector using the restriction enzymes *Bgl*II and *Not*I (NEB). A flexible linker of Gly(G)-Ser(S) of the sequence G-G-S-G-G-G-S-G-G-G-S-G-G was introduced between the LNS2 domain and GFP to allow independent and efficient folding of the two proteins. For cloning of the DDHD domain, the cDNA of RDGB corresponding to amino acids 730-913 was subcloned in *Bgl*II and *Xho*I digested pUAST-attB-mCherry with the flexible linker sequence present between mCherry and the DDHD domain. The DDHD-LNS2 construct was cloned by amplifying the cDNA corresponding to the amino acids 730-1259 of RDGB and tagging it to mCherry in *Bgl*II and *Xho*I digested pUAST-attB-mCherry, with the flexible linker sequence present between the mCherry and the DDHD domain

### Cell culture, transfection and immunofluorescence

S2R+ cells were cultured in Schneider’s insect medium (HiMedia) supplemented with 10% Fetal Bovine Serum and with antibiotics Penicillin and Streptomycin. Cells were transfected using Effectene (Qiagen) as per manufacturer’s protocol. Post 24 hours of transfection, cells were fixed with 4% paraformaldehyde (Electron Microscopy Sciences) and imaged to observe for GFP or mCherry fluorescence using a 60X 1.4 NA objective, in Olympus FV 3000 microscope

### Western Blotting

Heads of one day old flies were homogenised in 2X Laemmli sample buffer, and boiled at 95°C for 5 minutes. The samples were then run on a SDS-PAGE gel, and transferred on to a nitrocellulose membrane [Hybond-C Extra; (GE Healthcare, Buckinghamshire, UK)], with the help of a semi-dry transfer apparatus (BioRad, California, USA). The membrane was then blocked using 5% Blotto (sc-2325, Santa Cruz Biotechnology, Texas, USA) in Phosphate-buffered saline (PBS) with 0.1% Tween 20 (Sigma Aldrich) (PBST) for 2 hrs at room temperature (RT). The membrane was then incubated with the respective primary antibody, overnight at 4°C, using the appropriate dilutions [anti-RDGB (lab generated), 1:4000; anti-dVAP-A (kind gift from Dr. Girish Ratnaparkhi, IISER Pune), 1:3000; anti-α-tubulin-E7 (DSHB, Iowa, USA), 1:4000; anti-syntaxinA-8C3 (DSHB, Iowa, USA), 1:1000; anti-GFP (sc-9996), 1:2000]. Following this, the membrane was washed in PBST thrice, and incubated with the appropriate secondary antibody (Jackson Immunochemicals; dilution used: 1:10,000) coupled to horseradish peroxidase, at RT for 2 hrs. The blots were visualized using ECL (GE Healthcare), and imaged in a LAS4000 instrument.

### Immunostaining

For immunohistochemistry, retinae of one-day old flies were dissected under bright light in PBS. The samples were then fixed using 4% paraformaldehyde (Electron Microscopy Sciences) in PBS with 1 mg/ml saponin (Sigma Aldrich) for 30 minutes at RT. Post fixation, samples were washed thrice with PBS having 0.3% Triton X-100 (PBTX) and blocked using 5% Fetal Bovine Serum (ThermoFisher Scientific) in PBTX for 2 hrs at RT. The samples were then incubated overnight with the appropriate antibody in blocking solution at 4°C [anti-RDGB, (1:300); anti-GFP (1:5000), ab13970 (Abcam Cambridge, UK)]. Samples were then washed thrice with PBTX and incubated with the secondary antibody [Alexa Fluor 633 anti-rat (A21094), anti-chick (A21103), IgG (Molecular Probes)] at 1:300 dilution for 4 hrs at RT. For staining of the F-actin, Alexa Fluor 568–Phalloidin (Invitrogen, A12380) at 1:200 dilution was added during incubation with the secondary antibody. Samples were then washed in PBTX thrice and mounted with 70% glycerol in PBS. The whole-mounted preparations were imaged under 60X 1.4 NA objective, in Olympus FV 3000 microscope.

### Co-immunoprecipitation

S2R+ cells were co-transfected with mCherry::DDHD and LNS2::GFP for 48 hours, and lysed in ice-cold Protein Lysis Buffer [50mM Tris-Cl, 1mM EGTA, 1mM EDTA, 1% Triton X-100, 50mM NaF, 0.27 M Sucrose, 0.1% β-Mercaptoethanol]. 10% of the lysate was aliquoted to be used as input. The remaining lysate was split into two equal parts. To one part, anti-mCherry antibody (Thermo Fisher Scientific PA5-34974), (1.6 ug)] was added, and to the other part, a corresponding amount of control IgG (CST, 2729S) was added, and incubated overnight at 4°C. On the next day, Protein-G sepharose beads (GE Healthcare) were spun at 13000X g for 1 minute, and then washed with Tris-buffered saline (TBS), twice. The beads were then incubated with 5% Bovine Serum Albumin (BSA) (HiMedia) in TBS with 0.1%Tween-20 (TBST) for 2 hrs at 4°C. Equal amounts of blocked beads were then added to each sample, and incubated at 4°C for another 2 hrs. The immunoprecipitates were then washed twice with TBST containing β-Mercaptoethanol, and 0.1 mM EGTA for 5 minutes. The supernatant was then removed, and the beads were boiled in 2X Laemmli sample buffer for western blotting.

### Sub-cellular fractionation assay

The assay was performed as described by Sanxaridis et al., 2007 with minor modifications (Sanxaridis *et al*., 2007). Briefly, snap-frozen *Drosophila* heads were homogenised in ice-cold homogenisation buffer A (30 mM NaCl, 20 mM HEPES, 5 mM EDTA, pH=7.5). 10% of homogenate, representing the total head lysate, was directly taken for western blotting. The remaining homogenate was centrifuged at 5000 rpm for 5 minutes at 4°C to remove all chitinous material. The pellet was re-homogenized in the buffer to redeem any remaining membranous component from the cell ghost. This was done twice, post which the homogenate was spun at 100,000X g for 30 minutes, at 4°C to separate the entire membranous component from the cytosolic fraction. The pellet was reconstituted in buffer A. The re-suspended pellet representing the membrane fraction, and the supernatant representing the cytosolic fraction, were then individually used for doing western blotting.

### Lipid overlay assay

S2R+ cells were transfected with pJFRC-LNS2::GFP and pJRFC::GFP for 48 hours, following which cells were lysed with Protein Lysis Buffer (50 mM Tris-Cl, 1 mM EGTA, 1 mM EDTA, 1% Triton X-100, 50 mM NaF, 0.27 M Sucrose, 0.1% β-Mercaptoethanol). Commercially available PIP strips (Echelon Biosciences, P-6001) were blocked using 5% BSA (HiMedia) in TBST for 2 hours at RT. Following this, the strips were incubated overnight at 4°C with the remaining cell lysate. Next the membranes were washed extensively 5 times with 0.1% TBST and then incubated with anti-GFP antibody [(sc-9996), 1:2000] at RT for 2 hours. The membranes were then probed with HRP-conjugated anti-mouse IgG (Jackson Immunochemicals; 1:10,000) and binding was detected using ECL (GE Healthcare) in a LAS4000 instrument.

### Electrophysiology

Anaesthetised flies were immobilized at the end of a pipette tip by applying a drop of colourless nail polish on the proboscis. For recordings, GC 100F-10 borosilicate glass capillaries (640786, Harvard Apparatus, MA) were pulled to form electrodes and then filled with 0.8% (w/v) NaCl. The reference electrode was placed on the centre of the eye and the ground electrode on the thorax to obtain voltage changes post stimulation. The protocol for recording involved dark adapting the flies for 5 minutes initially, following which they were shown green flashes of light for 2 secs (10 times), and 12 secs of recovery time in dark between the two flashes. Voltage changes obtained were amplified using DAM50 amplifier (SYS-DAM50, WPI, FL), and recorded using pCLAMP10.7. Analysis was done using Clampfit 10.7 (Molecular Devices, CA). For analysis, the average of 10 recordings was taken for per fly.

### Deep pseudopupil imaging

The imaging is done with flies expressing a single copy of PH-PLCδ::GFP (PH domain of PLCδ, a PIP_2_ biosensor, tagged to GFP) driven by the transient receptor (*trp*) promoter of flies. Flies were anaesthetised and immobilized at the end of a pipette tip using a drop of colourless nail polish. The flies were then placed on the stage of an Olympus IX71 microscope, and the fluorescent pseudopupil focussed using a 10X objective lens. For imaging the deep pseudopupil, the flies were first adapted to red light for 6 minutes, following which a blue flash of 90 msec was given. The emitted fluorescence was captured, and its intensity was measured using Image J from NIH (Bethesda, Maryland, USA). Quantification of the fluorescence intensity was done by measuring the intensity values per unit area of the pseudopupil. The values are represented as mean +/-s.e.m.

## Figure Legends

**Supplemental Data 1:**

A. Western blot of protein extracts made from fly heads of RDGB^PITPd-FFAT^ and relevant controls. The blot is probed with antibody to RDGB. Syntaxin A1 is used as a loading control (N=3).

B. Western blot of protein extracts made from fly heads of RDGB^PITPd-FFAT^ and relevant controls expressing PH-PLCδ::GFP probe. The blot is probed with antibody to GFP. Syntaxin A1 is used as a loading control (N=3).

**Supplemental Data 2:**

A. Western blot of protein extracts made from fly heads of RDGB^(DDHD-LNS2)Δ^ and relevant controls. The blot is probed with antibody to RDGB. Syntaxin A1 is used as a loading control (N=3).

B. Representative ERG trace of 1 day old flies expressing RDGB^(DDHD-LNS2)Δ^ and relevant controls. Y-axis represents amplitude in mV, X-axis represents time in sec.

C. Representative images of fluorescent deep pseudopupil from 1 day old flies of expressing RDGB^(DDHD-LNS2)Δ^ and relevant controls expressing the PH-PLCδ::GFP probe.

D. Western blot of protein extracts made from fly heads expressing RDGB^(DDHD-LNS2)Δ^ and relevant controls and the PH-PLCδ::GFP probe. The blot is probed with antibody to GFP. Syntaxin is used as a loading control (N=3).

E. Western blot of protein extracts made from fly heads of RDGB^LNS2Δ^ and relevant controls. The blot is probed with antibody to RDGB. Tubulin is used as a loading control (N=3).

F. Representative ERG trace of 1 day old flies expressing RDGB^LNS2Δ^ and relevant controls. Y-axis represents amplitude in mV, X-axis represents time in sec.

G. Representative images of fluorescent deep pseudopupil from 1 day old flies of expressing RDGB^LNS2Δ^ and relevant controls along with the PH-PLCδ::GFP probe.

H. Western blot of protein extracts made from fly heads of RDGB^LNS2Δ^ and relevant controls, expressing PH-PLCδ::GFP probe. The blot is probed with antibody to GFP. Syntaxin A1 is used as a loading control (N=3).

**Supplemental Data 3:**

A. Western blot of protein extracts made from fly heads of RDGB^DDHD/4A^ and relevant controls. The blot is probed with antibody to RDGB. Tubulin is used as a loading control (N=3).

B. Western blot of protein extracts made from fly heads of RDGB^DDHD/4A^ and relevant controls expressing PH-PLCδ::GFP probe. The blot is probed with antibody to GFP. Syntaxin A1 is used as a loading control (N=3).

**Supplemental Data 4:**

A. PIP-Strip membranes were incubated over night with S2R+ cell lysate expressing LNS2::GFP and GFP for control. Binding is detected by probing with anti-GFP antibody [LPA= Lysophosphatidic acid, LPC= Lysophosphatidylcholine, PI= Phosphatidylinositol, PI3P= Phosphatidylinositol 3-phosphate, PI4P= Phosphatidylinositol 4-phosphate, PI5P= Phosphatidylinositol 5-phosphate, PE= Phosphatidylethanolamine, PC= Phosphatidylcholine, PS=Phosphatidylserine, PA= Phosphatidic acid, PI(3,4,5)P_3_= Phosphatidylinositol (3,4,5)-trisphosphate, PI(4,5)P_2_= Phosphatidylinositol (4,5)-bisphosphate, PI(3,5)P_2_=Phosphatidylinositol (3,5)-bisphosphate, PI(3,4)P_2_= Phosphatidylinositol (3,4) bisphosphate, S1P= Sphingosine-1-phosphate] (N=2 blots for LNS::GFP).

## References

Alli-Balogun, G. O. and Levine, T. P. (2019) ‘Regulation of targeting determinants in interorganelle communication.’, Current opinion in cell biology, 57, pp. 106–114. doi: 10.1016/j.ceb.2018.12.010.

Carvou, N. et al. (2010) ‘Phosphatidylinositol and phosphatidylcholine-transfer activity of PITP b is essential for COPI-mediated retrograde transport from the Golgi to the endoplasmic reticulum’, J.Cell.Sci, 15;123(Pt, pp. 1262–1273. doi: 10.1242/jcs.061986.

Chakrabarti, P. et al. (2015) ‘A dPIP5K dependent pool of phosphatidylinositol 4,5 bisphosphate (PIP2) is required for G-protein coupled signal transduction in Drosophila photoreceptors.’, PLoS genetics, 11(1), p. e1004948. doi: 10.1371/journal.pgen.1004948.

Chen, Y.-J., Quintanilla, C. G. and Liou, J. (2019) ‘Recent insights into mammalian ER–PM junctions’, Current Opinion in Cell Biology, 57, pp. 99–105. doi: 10.1016/j.ceb.2018.12.011.

Cockcroft, S. and Raghu, P. (2016) ‘Topological organisation of the phosphatidylinositol 4,5-bisphosphate-phospholipase C resynthesis cycle: PITPs bridge the ER-PM gap’, Biochemical Journal, 473(23), pp. 4289–4310. doi: 10.1042/BCJ20160514C.

Cockcroft, S. and Raghu, P. (2018) ‘Phospholipid transport protein function at organelle contact sites’, Current Opinion in Cell Biology, 53, pp. 52–60. doi: 10.1016/j.ceb.2018.04.011.

Cohen, S., Valm, A. M. and Lippincott-Schwartz, J. (2018) ‘Interacting organelles’, Current Opinion in Cell Biology, 53, pp. 84–91. doi: 10.1016/j.ceb.2018.06.003.

Dickeson, S. K. et al. (1989) ‘Isolation and sequence of cDNA clones encoding rat phosphatidylinositol transfer protein’, 264(28), pp. 16557–16564.

Gatta, A. T. and Levine, T. P. (2017) ‘Piecing Together the Patchwork of Contact Sites’, Trends in Cell Biology, 27(3), pp. 214–229. doi: 10.1016/j.tcb.2016.08.010.

Higgs, H. N. and Glomset, J. A. (1994) ‘Identification of a phosphatidic acid-preferring phospholipase A1 from bovine brain and testis’, Proc Natl Acad Sci USA., 91(20), pp. s9574–9578. doi: 10.1073/pnas.91.20.9574.

Inoue, H. et al. (2012) ‘Roles of SAM and DDHD domains in mammalian intracellular phospholipase A1 KIAA0725p.’, Biochimica et biophysica acta, 1823(4), pp. 930–9. doi: 10.1016/j.bbamcr.2012.02.002.

Kim, S. et al. (2013) ‘The phosphatidylinositol-transfer protein Nir2 binds phosphatidic acid and positively regulates phosphoinositide signalling.’, EMBO reports. Nature Publishing Group, 14(10), pp. 891–9. doi: 10.1038/embor.2013.113.

Kim, Y. J. et al. (2015) ‘Phosphatidylinositol-Phosphatidic Acid Exchange by Nir2 at ER-PM Contact Sites Maintains Phosphoinositide Signaling Competence.’, Developmental cell, 33(5), pp. 549–61. doi: 10.1016/j.devcel.2015.04.028.

Klinkenberg, D. et al. (2014) ‘A cascade of ER exit site assembly that is regulated by p125A and lipid signals’, Journal of Cell Science. Company of Biologists Ltd, 127(8), pp. 1765–1778. doi: 10.1242/jcs.138784.

Lev, S. et al. (1999) ‘Identification of a novel family of targets of PYK2 related to Drosophila retinal degeneration B (rdgB) protein’, 19(3), pp. 2278–88.

Nicita, F. et al. (2019) ‘Defining the clinical-genetic and neuroradiological features in SPG54: description of eight additional cases and nine novel DDHD2 variants’, Journal of Neurology. Dr. Dietrich Steinkopff Verlag GmbH and Co. KG, 266(11), pp. 2657–2664. doi: 10.1007/s00415-019-09466-y.

Pensato, V. et al. (2014) ‘Overlapping phenotypes in complex spastic paraplegias SPG11, SPG15, SPG35 and SPG48’, Brain. Oxford University Press, 137(7), pp. 1907–1920. doi: 10.1093/brain/awu121.

Raghu, P., Yadav, S. and Mallampati, N. B. N. (2012) ‘Lipid signaling in Drosophila photoreceptors.’, Biochimica et biophysica acta, 1821(8), pp. 1154–65. doi: 10.1016/j.bbalip.2012.03.008.

Saheki, Y. and Camilli P. De (2017) ‘Endoplasmic Reticulum – Plasma Membrane Contact Sites’, Annual Review of Biochemistry, 86, pp. 659–684.

Sanxaridis, P. D. et al. (2007) ‘Light-induced recruitment of INAD-signaling complexes to detergent-resistant lipid rafts in Drosophila photoreceptors.’, Molecular and cellular neurosciences, 36(1), pp. 36–46. doi: 10.1016/j.mcn.2007.05.006.

Tesson, C. et al. (2012) ‘Alteration of fatty-acid-metabolizing enzymes affects mitochondrial form and function in hereditary spastic paraplegia’, American Journal of Human Genetics. Am J Hum Genet, 91(6), pp. 1051–1064. doi: 10.1016/j.ajhg.2012.11.001.

Vihtelic, T. S. et al. (1993) ‘Localization of Drosophila retinal degeneration B, a membrane-associated phosphatidylinositol transfer protein.’, The Journal of cell biology, 122(5), pp. 1013–22.

Yadav, S. et al. (2015) ‘RDGBα, a PtdIns-PtdOH transfer protein, regulates G-proteincoupled PtdIns(4,5)P2 signalling during Drosophila phototransduction’, Journal of Cell Science, 128(17), pp. 3330–3344. doi: 10.1242/jcs.173476.

Yadav, S. et al. (2018) ‘RDGBα localization and function at a membrane contact site is regulated by FFAT/VAP interactions.’, Journal of Cell Science, 131(ojcs207985), p. doi:10.1242/jcs.207985. doi: 10.1242/jcs.207985.

Yadav, S., Cockcroft, S. and Raghu, P. (2016) ‘The Drosophila photoreceptor as a model system for studying signalling at membrane contact sites.’, Biochemical Society transactions, 44(2), pp. 447–51. doi: 10.1042/BST20150256.

