## Supplementary figures and images for "Interdomain interactions regulate the localization of a lipid transfer protein at ER-PM contact sites"

### Supplemental Data

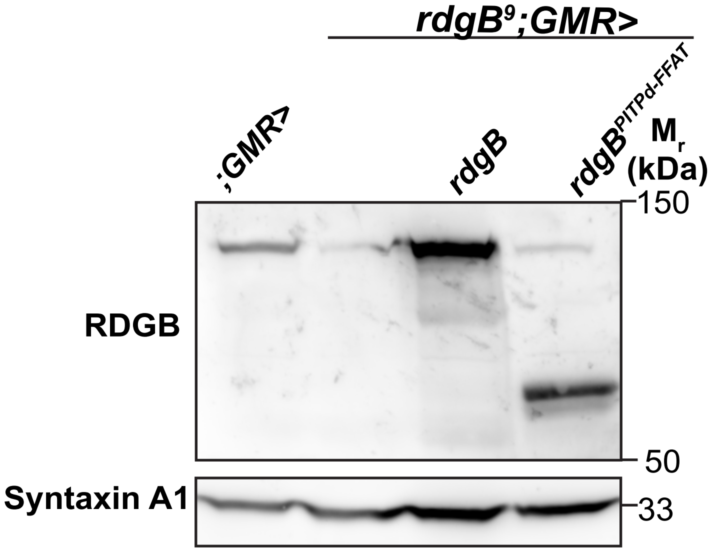

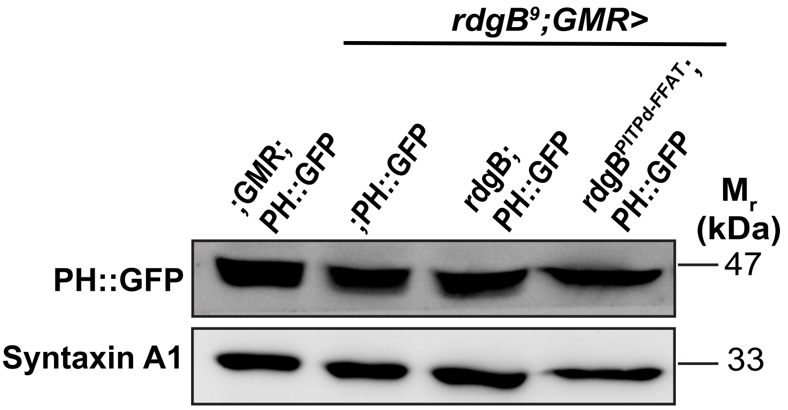

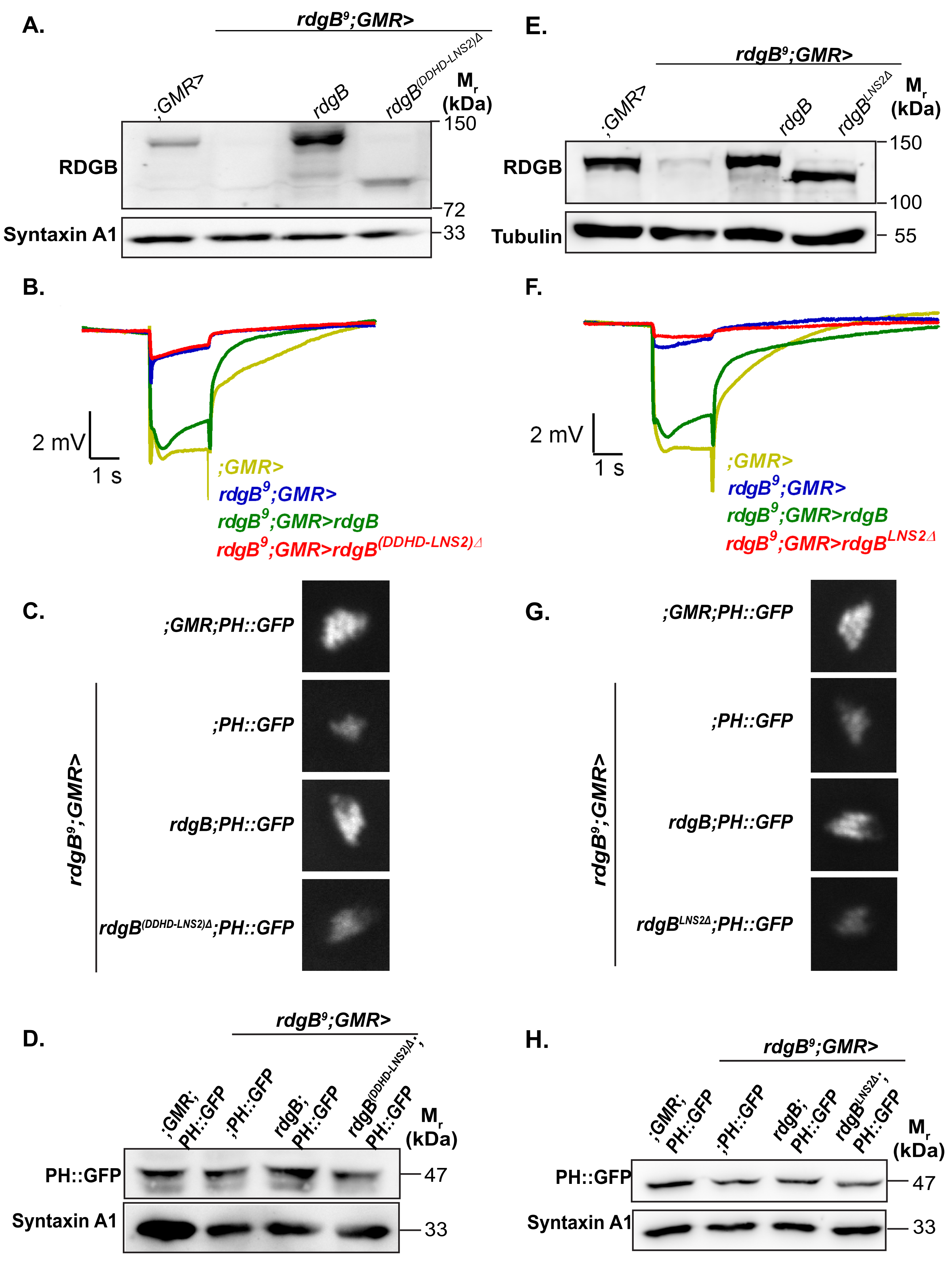


Basak et al., Supplemental Data 2

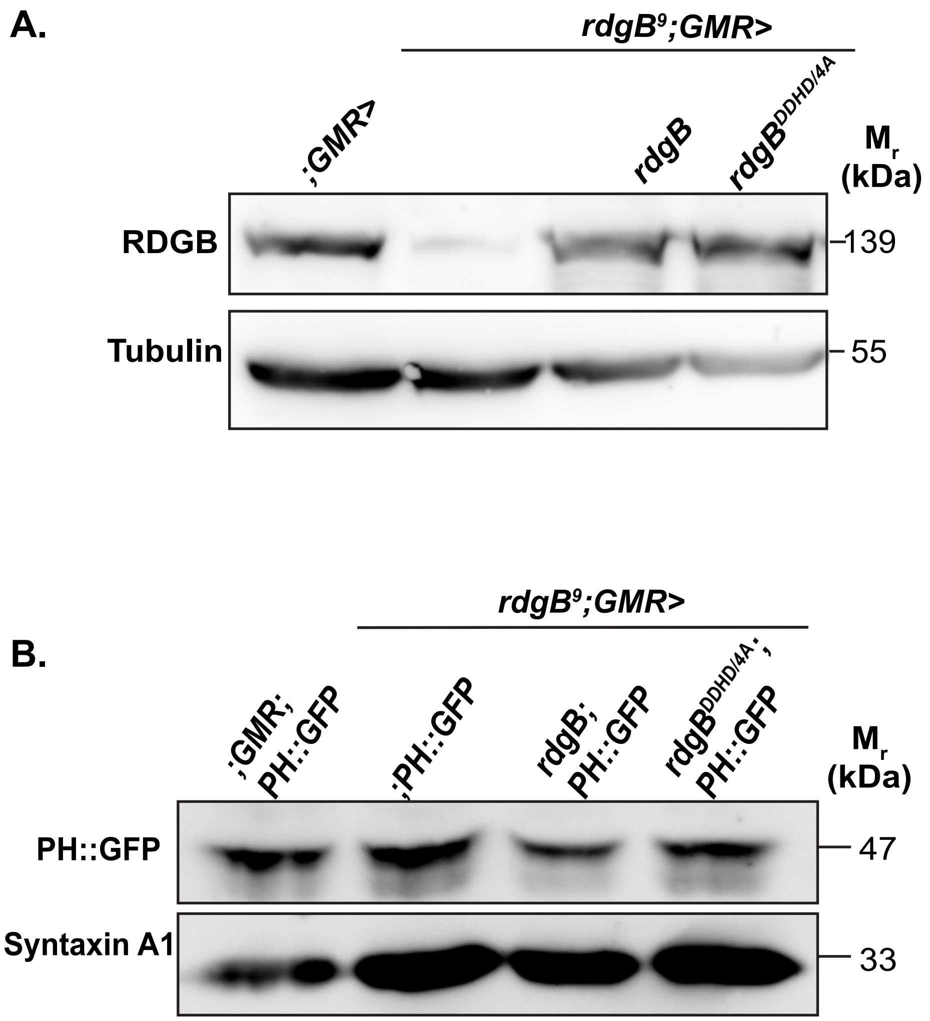

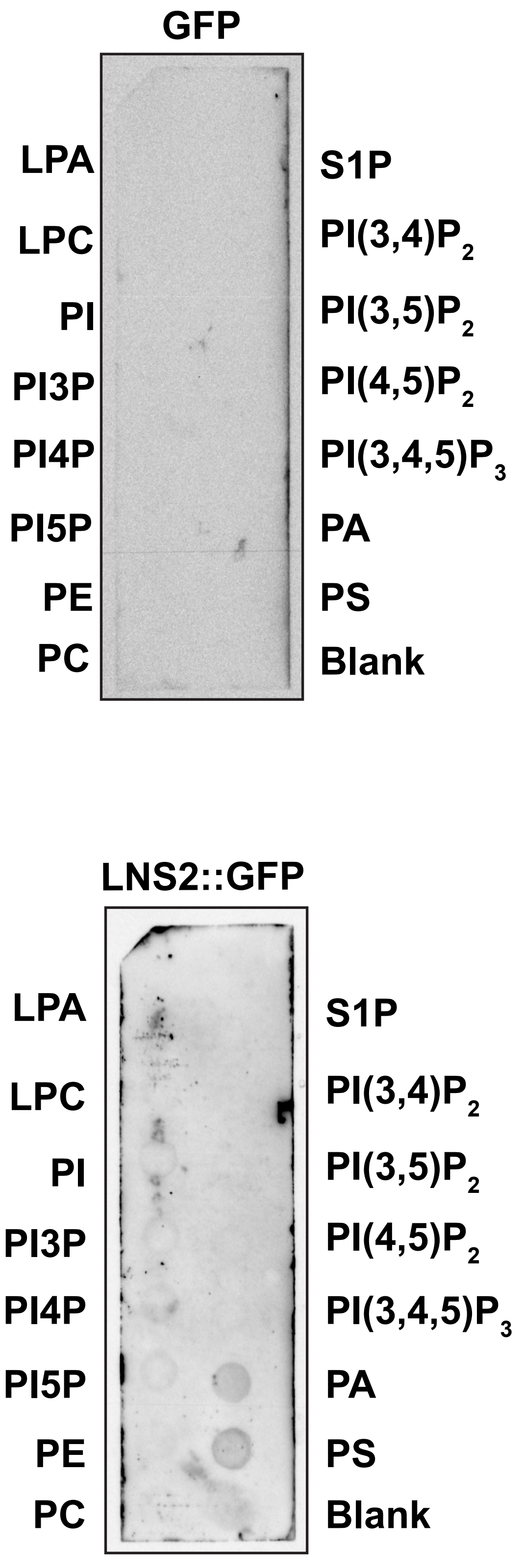
